# ctmmweb: A graphical user interface for autocorrelation-informed home range estimation

**DOI:** 10.1101/2020.05.11.087932

**Authors:** Justin M. Calabrese, Christen H. Fleming, Michael J. Noonan, Xianghui Dong

## Abstract

Estimating animal home ranges is a primary purpose of collecting tracking data. All conventional home range estimators in widespread usage, including minimum convex polygons and kernel density estimators, assume independently sampled data. In stark contrast, modern GPS animal tracking datasets are almost always strongly autocorrelated. This incongruence between estimator assumptions and empirical reality leads to systematically underestimated home ranges. Autocorrelated kernel density estimation (AKDE) resolves this conflict by modeling the observed autocorrelation structure of tracking data during home range estimation, and has been shown to perform accurately across a broad range of tracking datasets. However, compared to conventional estimators, AKDE requires additional modeling steps and has heretofore only been accessible via the command-line `ctmm` R package. Here, we introduce `ctmmweb`, which provides a point-and-click graphical interface to `ctmm`, and streamlines AKDE, its prerequisite autocorrelation modeling steps, and a number of additional movement analyses. We demonstrate `ctmmweb’s` capabilities, including AKDE home range estimation and subsequent home range overlap analysis, on a dataset of four jaguars from the Brazilian Pantanal. We intend `ctmmweb` to open AKDE and related autocorrelation-explicit analyses to a wider audience of wildlife and conservation professionals.

---

Home range estimation is one of the most routinely applied analyses to animal tracking data. Though many home range estimators exist, all methods in common use, including minimum convex polygons (MCPs; Mohr 1947), local convex hulls (LoCoHs; Getz et al. 2007), and conventional kernel density estimators (KDEs; Worton 1989), assume independently sampled data (Laver and Kelly, 2008). In stark contrast, modern GPS tracking datasets are nearly always autocorrelated (Noonan et al. 2019; Noonan et al. 2020). Regardless of their means of locomotion (e.g., walking, hopping, flying, swimming, etc.), animals trace continuous paths through the environment (Turchin 1997). An unavoidable consequence of the continuity of animal movement paths is that finer sampling in time leads to more strongly autocorrelated tracking data, all else being equal (Swithart and Slade 1985). GPS sampling rates have increased in lock step with technological advances, and it is becoming fairly common to speak of sampling rates in terms of hertz (observations per second; Kays et al. 2015, Noonan et al. 2015). Looking to the future, the strength of autocorrelation in tracking data will likely continue to increase as improvements in tracking technology facilitate the ever-finer sampling of animal paths.

There has been a long-running debate about how problematic (or not) autocorrelated data are for home range estimators that assume Independently and Identically Distributed (IID) data (Swithart and Slade 1985, de Solla et al. 1999, Blundell et al. 2001, Fieberg 2007). Conclusions in this literature, which are typically based on small datasets and limited ranges of observed autocorrelation, span the gamut from “autocorrelation is a serious problem” to “autocorrelation is a red herring” to “autocorrelation is actually beneficial”. Recent work based on an unprecedented empirical dataset (369 individuals, 27 species, five continents) covering the full range of autocorrelation strengths, has shown conclusively that, when applied to autocorrelated data, IID home range estimators are negatively biased (Noonan et al. 2019). Furthermore, the magnitude of the bias increases in proportion to the strength of autocorrelation in the data (Noonan et al. 2019). Consequently, addressing the issue of autocorrelation in animal tracking data is a pre-requisite for accurate home range estimation. Only two viable solutions to this problem exist: either the data must be thinned to reduce autocorrelation and more closely approximate the IID assumption, which is wasteful, inefficient, and defeats the purpose of collecting finely-sampled data, or the autocorrelation in the data must be appropriately modeled (Fleming et al. 2015).

A further requirement for accurately quantifying home ranges is using an estimator of the range distribution, as opposed to an estimator of the occurrence distribution (Fleming et al. 2015, Fleming et al. 2016, Horne et al. 2020). The range distribution quantifies the animal’s long-run space use and is consistent with Burt’s (1943) conceptual definition of home range. In contrast, the occurrence distribution quantifies uncertainty in the animal’s true location during the tracking period, and thus collapses to the animal’s movement path as data quality improves and uncertainty in its location disappears. This means that the occurrence distribution’s area shrinks towards zero as data become more finely and accurately sampled. Conventional KDEs and MCPs belong to the family of range estimators, but do not model autocorrelation in the data. Conversely, the widely used Brownian Bridge (Horne et al. 2007) accounts for autocorrelation in the data, but belongs to the family of occurrence esitmators. Therefore, none of these approaches solve the problem of home range estimation with autocorrelated data.

Autocorrelated Kernel Density Estimation (AKDE) is the first method that explicitly models the autocorrelation in the data, and belongs to the family of range distribution estimators (Fleming et al. 2015, Fleming and Calabrese 2017). The large-scale comparative analysis of Noonan et al. (2019) showed conclusively that AKDE is currently the only proper range estimator that is consistently accurate, as evidenced by block cross-validation performance (Roberts et al. 2017), across the full spectrum of sample sizes and autocorrelation strengths represented in their extensive dataset. In contrast, IID estimators on average underestimated home range areas by a factor of two (MCP and Gaussian reference function KDE) or a factor of 13 (k-LoCoH and least squares cross-validation KDE) across their dataset (Noonan et al. 2019). For individuals plagued by small effective sample size (i.e., few range crossings), IID estimates showed an even larger degree of underestimation (Noonan et al. 2019).

While the benefit of explicitly modeling autocorrelation in the data is improved accuracy (Fleming and Calabrese 2017, Winner et al. 2018, Noonan et al. 2019), the cost is greater analytical complexity and extra modeling steps that must be taken prior to home range estimation (Fleming et al. 2015, Calabrese et al. 2016, Fleming and Calabrese 2017, Fleming et al. 2019). Specifically, an appropriate model for the autocorrelation structure of the data must first be identified, and then used as a basis for AKDE home range estimation (Fleming et al. 2015). The R package `ctmm` implements these prerequisite modeling steps, as well as AKDE estimation and a suite of related analyses (Fleming and Calabrese 2015, Calabrese et al. 2016), but requires R programming knowledge. To make AKDE available to a broader audience, we introduce `ctmmweb` (Dong et al. 2018), which is an R Shiny-based (Cheng et al. 2019) graphical user interface to the `ctmm` package, and includes additional functionality for publication-quality graphics, interactive maps, and reproducible research. We demonstrate the capabilities of `ctmmweb`, and its workflow for home range and overlap analysis, on an example featuring jaguars (*Panthera onca*) in the Brazilian Pantanal (Morato et al. 2018).

## METHODS

While `ctmmweb` has a wide range of capabilities and can perform many types of movement analyses, in this introductory paper we detail only the steps necessary to reproduce the home range and overlap analysis. The app consists of a series of pages, with each page containing related functionality. For example, the Import page supports several different means of importing data into `ctmmweb`. Within each page, boxes are used to further separate different functions. Help buttons occur within boxes and provide guidance on the functionality contained in the focal box. The pages are indexed in `ctmmweb’s` sidebar on the left side (Fig. 1), and a typical analysis proceeds by moving down the sidebar sequentially from one page to the next. Below, we discuss each page required to estimate the home ranges and overlap of four jaguars in the Brazilian Pantanal.

**Figure 1.**
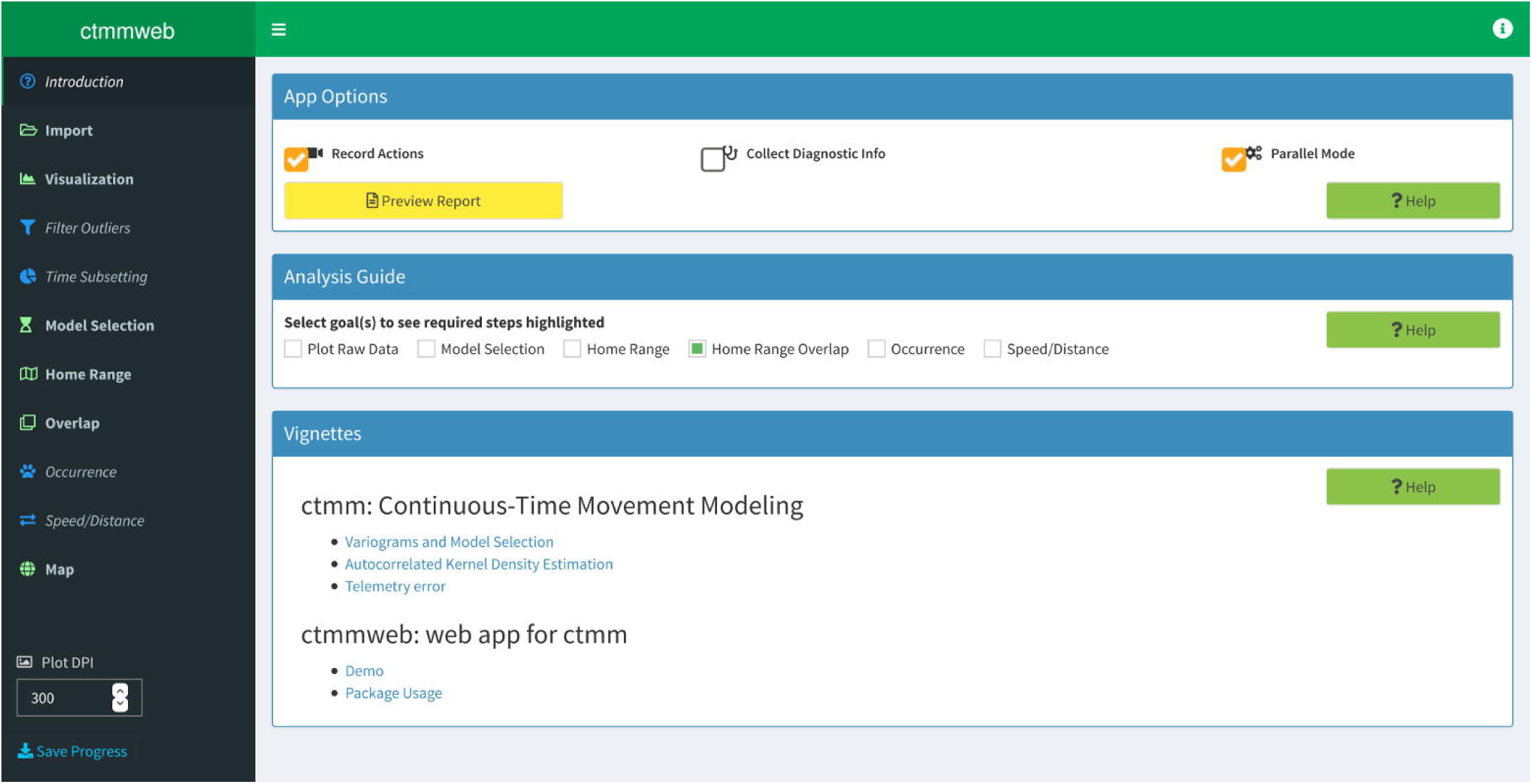
The user interface of `ctmmweb`. The analysis workflow proceeds from page to page down the left-hand sidebar, though some pages may be skipped depending on analysis goals. The Introduction page is currently displayed, showing the Analysis Guide feature, which in this case, highlights the workflow steps necessary to complete home range overlap analysis.

### Introduction

The introduction page allows the user to configure certain app settings (e.g., whether or not to use multiple processor cores for parallelization), and provides guidance on how to use `ctmmweb`. To facilitate reproducible research, the “Record Actions” checkbox, which is checked by default, ensures that the app records all actions, datasets used, analyses, results, and figures. An archive of the user’s `ctmmweb` session can be downloaded at any time (and from any page) by pressing the “Save Progress” button on the bottom of the sidebar. In addition to containing the aforementioned elements, the archive also provides a combined. html work report that can serve as a detailed description of the `ctmmweb` session. The “Analysis Guide” provides visual cues for the user to follow to accomplish an analysis goal or goals. The guide is off by default and requires the user to select a goal (or set of goals) from the listed checkboxes. The required steps to achieve the chosen goal are then show with boldface type and a green icon in the sidebar. The user then only needs to follow the highlighted steps sequentially from the top to the bottom of the sidebar to achieve their selected goal. Figure 1 shows the analysis guide for the goal of home range overlap analysis that we will demonstrate on the jaguar data. Finally, a series of vignettes on `ctmmweb` and the analyses it supports are provide to give more detailed guidance.

### Import

The app assumes that data are in Movebank.org format (Wikelski and Kays 2014), which is a tabular,. csv plain text format with one observation per row and columns that minimally include: *individual.local.identifier* (or *tag.local.identifier*), *timestamp*, *location.long* and *location.lat*. One can either manually create (e.g., in R or a spreadsheet program) an appropriately formatted. csv by including and correctly naming the above-listed elements, or use Movebank. For the latter option, one can either download a. csv from Movebank and manually import it into `ctmmweb` (via “Upload Data”), or use `ctmmweb’s` “Import from Movebank” dialog to directly import the data into the app without requiring the data be stored locally. It is also possible to import a previously saved `ctmmweb` session archive via the “Restore Progress” dialog, so that the researcher can pick up where they left off, perform additional analyses and/or modify existing ones, and save the results in an updated archive. The jaguar data used in this paper can be imported by selecting “jaguar” from the list of built-in datasets and clicking “load”.

### Visualization

The imported individuals are listed in a table on the Visualization page, which provides both descriptive statistics as well as a means for the user to select which individuals to include in the analyses. This selection of individuals carries on throughout the rest of the app. A number of basic graphics are available on the Visualization page. For example, a scatterplot shows the raw (*x*, *y*) locations of all selected individuals, color coded by individual, while facet plots show the (*x*, *y*) locations of each individual in its own panel, with the same axis ranges used across all panels to facilitate comparison. All of the figures produced in the app will be saved to the session archive upon clicking “Save Progress” at the resolution indicated in the “Plot DPI” box immediately above the save button.

### Model Selection

Variogram analysis, which is available on the Model Selection page, is the core visual diagnostic used to examine the autocorrelation structure of a tracking dataset (Fleming et al. 2014, Calabrese et al. 2016). Variograms depict the averaged squared displacement between all pairs of points separated by a given time lag (e.g., 1 hour), plotted as a function of time lag. This representation of the data allows one to visually assess, among other features, the appropriateness of home range analysis and the degree to which there is directional persistence in the movement. In the former case, an asymptote at larger time lags indicates evidence of range residency in the data, which implies home range analysis is appropriate. Conversely, the lack of a clear asymptote would indicate no evidence for range residency and would suggest that home range analysis is inappropriate for the focal dataset. While analyses other than home range estimation are still possible with non-range-resident data, `ctmmweb` does not currently support the non-range-resident movement models required in such cases. The user would instead have to turn to ctmm. Second, upward curvature on the short-lag end of the variogram is evidence of correlated velocity in the data, and suggests that a correlated velocity model would be appropriate (see model selection, below). Recalling that velocity consists of both movement speed and direction of travel, correlated velocity implies directional persistence in the movement. `ctmmweb` features rich functionality for variograms and by default produces a multi-panel display featuring a variogram for each individual in the dataset (Fig. 2). Superimposed on each variogram is an algorithmically generated “guess” at initial parameter values for fitting a movement model to the data. If the visual correspondence between the empirical variogram and the “guess” is poor, the underlying parameter values can be manually adjusted for each individual via a dialog box with parameter sliders.

**Figure 2.**
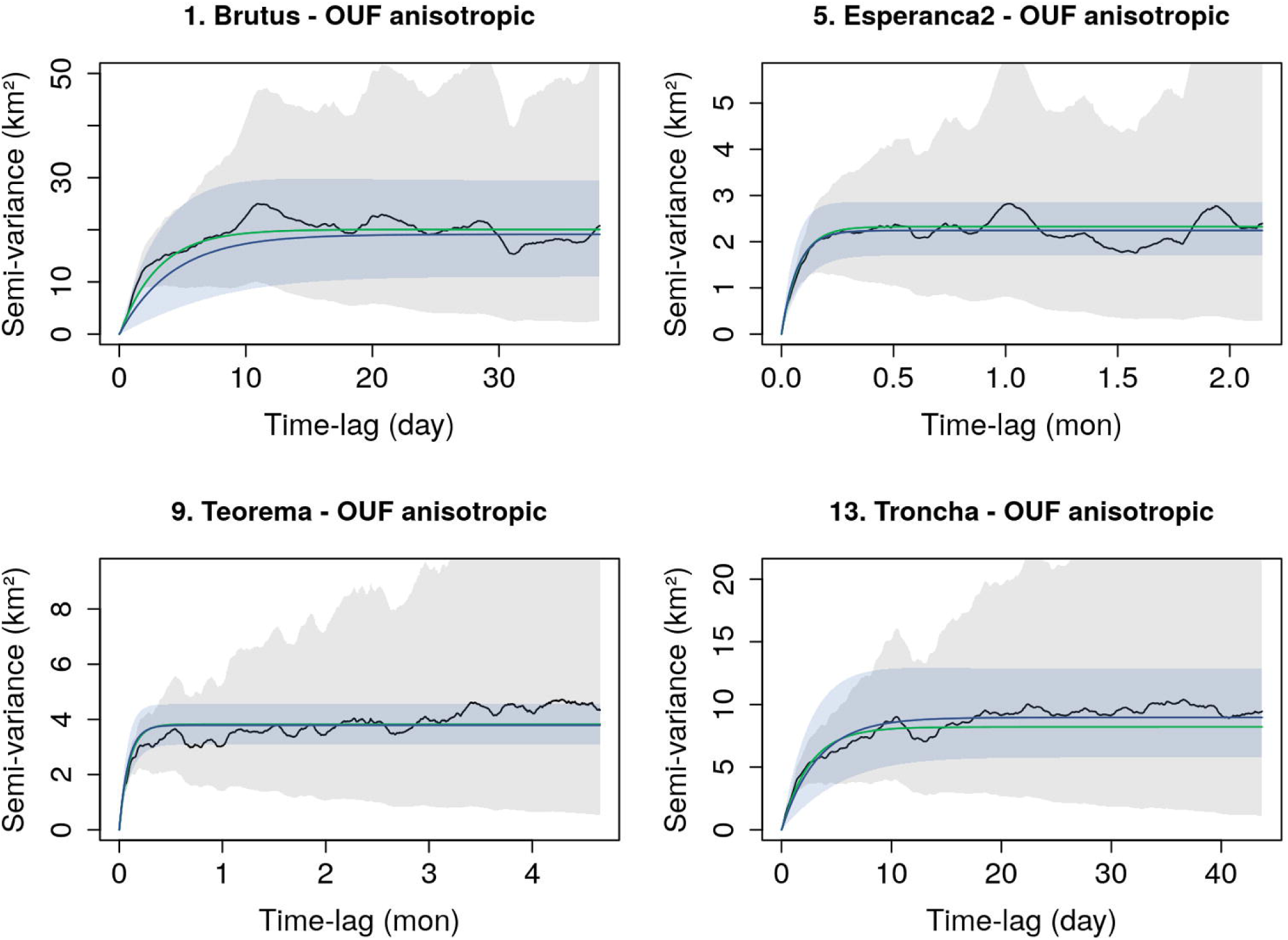
Empirical variograms (black curves and gray confidence envelopes) and semi-variance function of the AIC-best model (blue curves and blue confidence envelopes) for each individual. The green curves are the semi-variance functions implied by the intial parameter guesses. By default, `ctmmweb` shows the first 50% of each variogram, in keeping with standard practice in geostatistics. All four jaguars show strong evidence of range residency with each variogram having a clear asymptote, and the OUF-Anisotropic model was selected for all individuals.

After visual examination of the data, clicking the “Modeled” tab on the Model Selection page performs automated model fitting and selection, based on the initial parameter value guess identified above. Model fitting is done via perturbative Hybrid Residual Maximum Likelihood (pHREML; Fleming et al. 2019) estimation, and model selection is based on Akaike’s Information Criterion (AIC; Burnham and Anderson 2002, Fleming et al. 2019). For data with evidence of range residency, the set of candidate models includes the IID process, the Ornstein-Uhlenbeck (OU) process (Ornstein and Uhlenbeck 1930), and the OU-Foraging process (OUF; Fleming et al. 2014). The IID process, which is the model assumed by conventional range estimators such as KDE and MCP, has a home range but both positions and velocities are uncorrelated. The OU process features a home range, correlated positions, but uncorrelated velocities (i.e., no directional persistence). Finally, the OUF process is the most general model, and includes a home range, correlated positions, and correlated velocities. Each of these movement processes is considered both in isotropic form, meaning the movement is the same in all directions (i.e., circular home range), or anisotropic form (i.e., non-circular home range), where the movement may vary by direction, leading to a set of six core candidate models. After the fitting and model selection algorithms have run, the selected model is displayed graphically via the variogram for each individual, and the details of the fit are output in a table below the variograms. For each individual, the AIC-best model is highlighted by default in the results table, and will be used as the basis for the conditional analyses that follow. If desired (i.e., for comparative purposes), the user can choose to base downstream conditional analyses on any other model in the results table by manually selecting it. As a rule, however, optimal results will only be obtained by conditioning on the AIC-best model for each individual. The parameter estimates displayed in the table for each model may also be of interest. For example, the parameter τ[position] gives the average time it takes the focal individual to cross the linear extent of its home range. It is important to realize that because `ctmmweb` is considering many different autocorrelation models in the background (instead of assuming the model *a priori* as in conventional movement analyses), the model selection step is computationally intensive and can take substantial time for large datasets.

### Home Range

AKDE home range analysis can be based on any range-resident movement model supported by `ctmmweb`. Importantly, AKDE based on the IID model reduces exactly to conventional KDE with the Gaussian reference function bandwidth optimizer (Fleming et al. 2015). This means that for a dataset with statistically independent locations, `ctmmweb` will select the IID model as AIC-best, and a conventional KDE estimate will be produced. If instead the data are autocorrelated, the most appropriate autocorrelated movement model (either OU or OUF) will be selected and used as the basis for AKDE. Assuming a one-size-fits-all model *a priori*, like conventional home range estimators do via their IID assumption, can result in home ranges that are underestimated in proportion to the strength of autocorrelation in the data. This issue can be avoided by letting the data inform the choice of an appropriate autocorrelated movement model via model selection. After all, it is not autocorrelation *per se* that causes problems, but rather *unmodeled* autocorrelation and the resulting violation of the IID assumption made by conventional estimators.

After model fitting and selection has finished, selecting the Home Range page from the navigation panel will automatically calculate and plot AKDE estimates for each individual. The app also provides a table of summary statistics about the AKDE estimates, including the area corresponding to the focal percentile of the range distribution (95% by default, but user adjustable), as well as confidence intervals on this estimate.

### Overlap

As individuals, by definition, perform most of their movements within their home ranges, one can expect home range overlap to be proportional to encounter rates among individuals (Martinez-Garcia et al. 2020). Thus overlap can serve as a useful proxy for interaction potential. After home range estimation, pairwise overlap among individuals can be estimated via the Overlap page. In `ctmmweb`, overlap is quantified by the widely-used Bhattacharyya coefficient (Fieberg and Kochanny 2005, Winner et al. 2018), which is a symmetric index and ranges between 0 (no overlap) and 1 (complete overlap). When overlap analysis is the goal, AKDE home ranges must be calculated on the same grid, which is the default choice but comes at the expense of longer run times.

### Map

To help contextualize the results, `ctmmweb` allows both the data and home range estimates to be plotted on an interactive map provided by the R interface to the Java Leaflet library (Cheng et al. 2018). The user can select which individuals, and which elements for each individual (data and/or home range estimates), to display. Furthermore, the user can choose among a range of map options including terrain, satellite, and topographic backgrounds.

## STUDY AREA

To demonstrate the utility of `ctmmweb`, we use GPS data from four jaguars (one male and three females) tracked in the Brazilian Pantanal between 2013 and 2015 (Morato et al. 2018). With an estimated area of 150,355 km^2^, the Pantanal is the world’s largest tropical wetland, and is a biome in which Jaguars are considered to be vulnerable (Morato et al. 2016). Jaguars are medium sized (ca. 75 kg), range-resident carnivores that cross their home ranges on hourly timescales (Morato et al. 2016). For these individuals, locations were collected at hourly intervals for 141 days on average (range: 76 – 274 days) via Lotek/Iridium GPS collars, resulting in a mean of 2480 recorded locations per animal (range: 1323 - 4860).

## RESULTS

Using the jaguar data, we completed all steps of the workflow from import through conditional analyses including AKDE home range estimation and home range overlap via the Bhattacharyya coefficient. At the end of the analysis session, we saved our results using the “save progress” button at the bottom of `ctmmweb`’s side bar (Fig. 1). The full archive, including the work report, produced from this analysis session is provided in the Supporting Information S1.

All four jaguars show clear range residency, as evidenced by the pronounced asymptotes in their empirical variograms (Fig. 2). For all four individuals, the OUF-Anisotropic model was unequivocally selected, with AIC differences in favor of this model >8 in all cases, and >69 in two out of four cases (see the model selection results table in the work report in Supplementary Material S2 for full details). This model features correlated positions, correlated velocities (directional persistence), and a home range, and all of these features were apparent in the empirical variograms of the jaguars (Fig. 2). The “anisotropic” moniker on the selected model name implies that each animal’s movement was spatially asymmetrical, i.e., greater in some directions than in others.

We next computed AKDE home range estimates for all four individuals, which `ctmmweb` displays individually on the Home Range page (Fig. 3, top row). Notice the grid superimposed on the background of the density estimates. The size of this grid depicts the bandwidth size and represents the spatial resolution of the home range estimate. Features apparent in the home range (e.g., holes) that are smaller than this grid should be ignored because they are below the spatial resolution of the estimate and thus could be spurious. Said another way, the grid represents the finest spatial scale that can be accurately resolved by the home range estimate. Figure 3 also shows the data and home range estimates from all four animals overlaid on a topographical map, using `ctmmweb`’s interactive Map page. This view of the home range estimates suggests significant pairwise overlap among individuals, particularly for the lone male jaguar, Brutus, with the three females.

**Figure 3.**
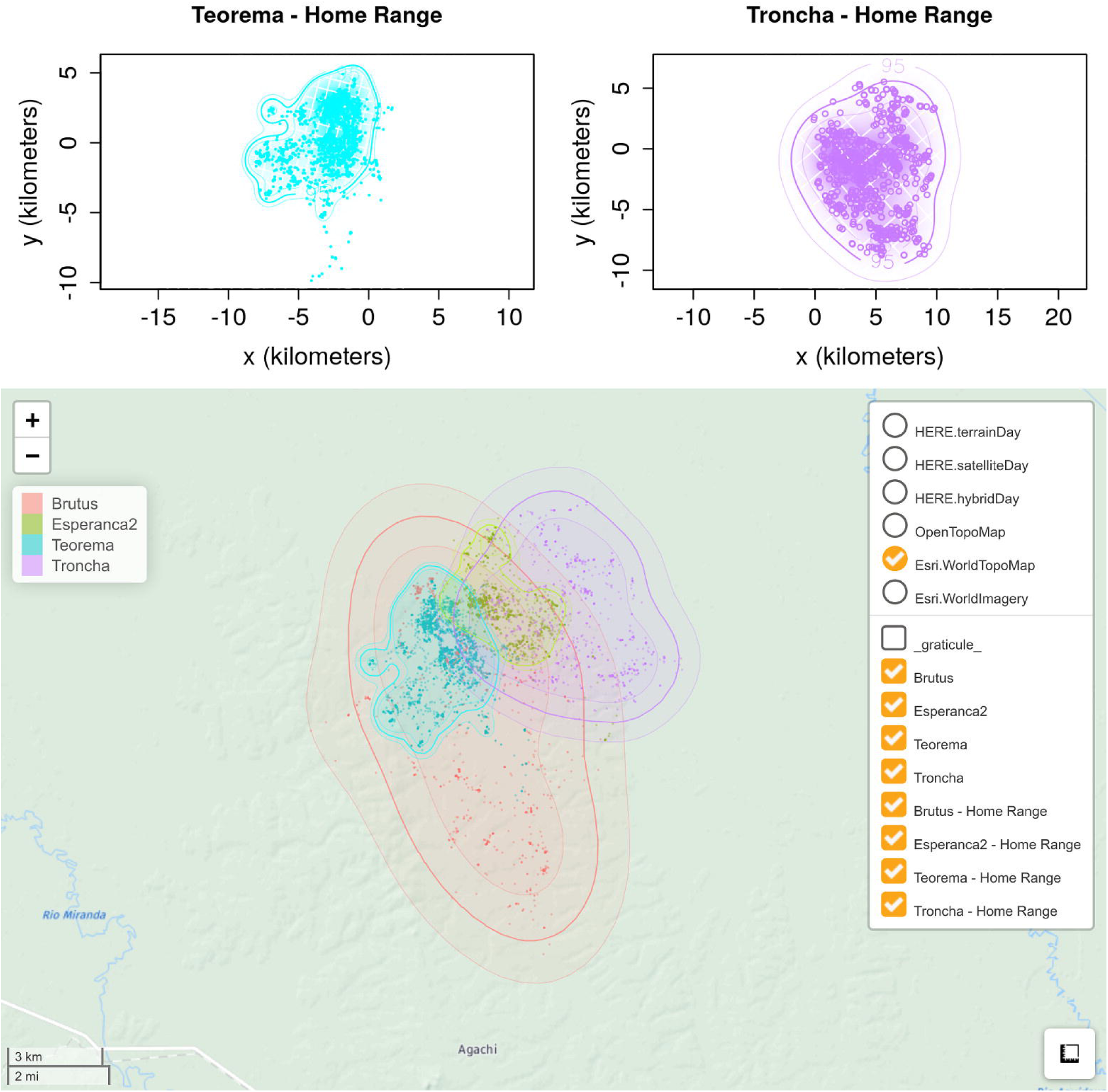
Top row: AKDE estimates (density + contours) and data (filled circles) for Teorema and Troncha, as they are displayed on the Home Range page of the app. Notice the bandwidth grid in the background of these two estimates. Bottom row: AKDE home range estimates, together with the location data, for all four individuals plotted on an interactive topological map via the Map page. All estimates display the 95% contour of the range distribution (thick curve) and the 95% confidence envelopes (thin curves) on the contour estimates.

To properly quantify this overlap, we calculated the Bhattacharyya coefficient via `ctmmweb`’s Overlap page, given the AKDE estimates previously calculated. A visual depiction of overlap for two pairs of individuals is shown in the top row of Figure 4, while the overlap estimates and 95% confidence intervals are shown for all pairs of individuals in the bottom panel of Figure 4. All pairwise overlaps were statistically significant (lower confidence limit not including zero overlap), but the average overlap of the lone male with the females (0.52 [0.41, 0.63]; point estimate with 95% confidence intervals) was significantly greater (p < 0.05) than average pairwise overlap among females (0.33 [0.25, 0.40]), with the estimated difference between M-F and F-F overlap being 0.20 [0.06, 0.33].

**Figure 4.**
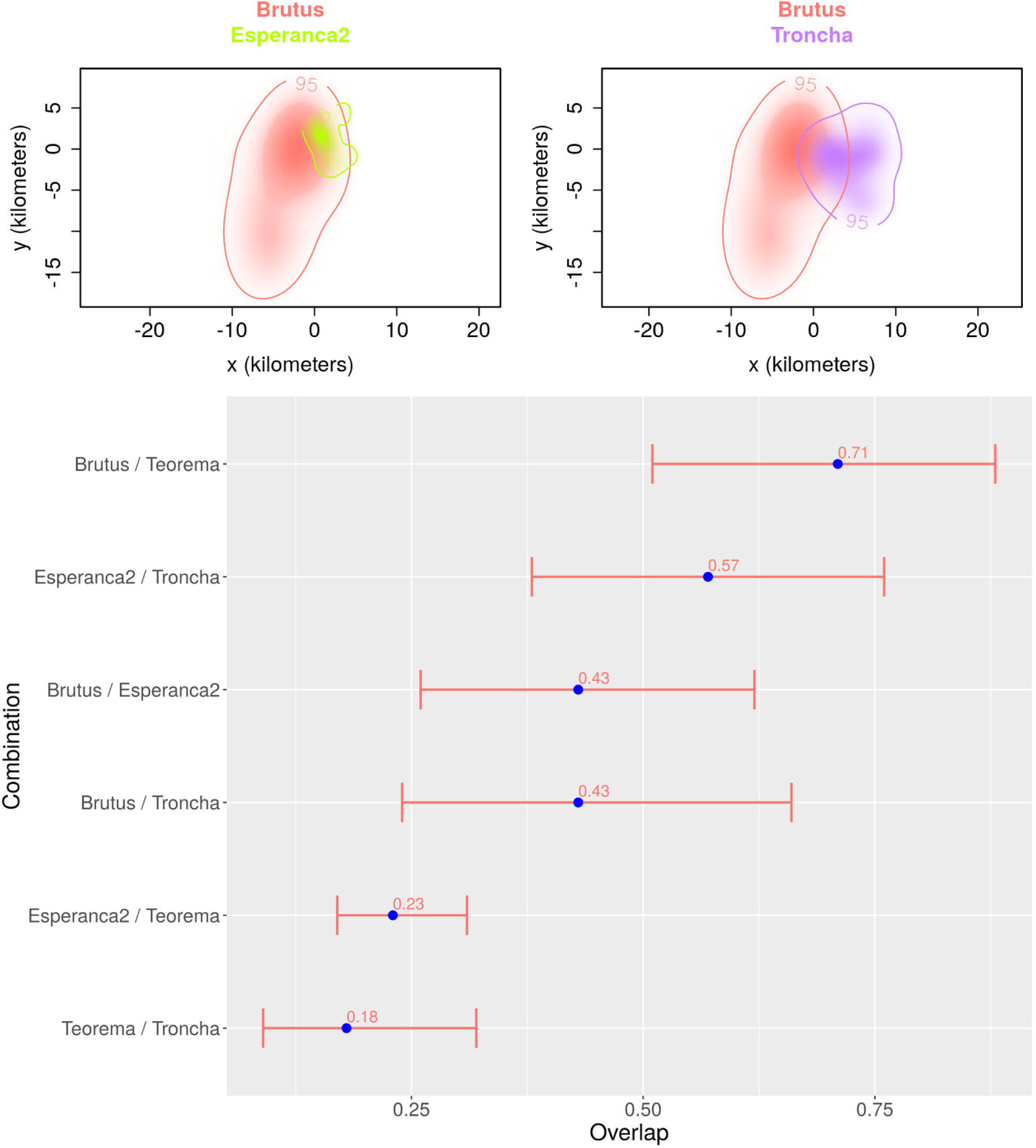
Top row: Visual representation of home range overlap based on the estimated AKDE densities for Brutus/Esperanca2 (left) and Brutus/Troncha (right). Bottow row: Pairwise overlap estimates based on the Bhattacharyya coefficient (blue points) and 95% confidence intervals (red bars) for all possible pairs of individuals.

Additional results and figures from the jaguar analysis can be found in the work report that is included in Supplementary Material S1.

## DISCUSSION

Autocorrelation has become a critical issue in animal tracking data that can no longer be ignored. Analyses based on such data will either have to account for autocorrelation or risk being systematically, and often grossly, biased. Fortunately, substantial progress in accounting for autocorrelation in animal tracking analyses has been made in recent years. The bulk of these advances, however, have heretofore only been accessible in command-line analysis packages such as `ctmm` (Fleming and Calabrese 2015, Calabrese et al. 2016), `crawl` (Johnson et al. 2008), `move` (Kranstauber and Smolla 2013), `corrMove` (Calabrese et al. 2018), and `marcher` (Gurarie et al. 2017). Of these, `ctmm` is by far the most comprehensive, and `ctmmweb` provides easy point-and-click access to the vast majority of analyses supported by ctmm, including AKDE home range estimation. An additional advantage of `ctmmweb`, is that all of the statistical estimates it produces are accompanied by accurate confidence intervals. To our knowledge, no other software packages besides `ctmmweb` and `ctmm` provide confidence intervals on home range and overlap estimates.

Focusing on data from four jaguars in Brazil, we have demonstrated how `ctmmweb` can be used to visualize and understand key features in tracking data, fit and select appropriate movement models, and perform conditional analyses such as AKDE home range estimation and home range overlap estimation. Home range estimates were very similar to those previously reported for these four jaguars by Morato et al. (2016), with minor differences accounted for by refinements made to the underlying `ctmm` package since Morato’s study. In contrast, the overlap estimates reported here for the jaguars are novel and demonstrate lower pairwise overlap of home ranges among neighboring females than among male-female pairs. This result is consistent with reports from the literature on both territoriality among females, and on male home ranges overlapping with multiple females (Schaller and Cranshaw 1980, Rabinowitz and Nottingham 1986). Furthermore, it is also the spatial arrangement expected under the jaguar’s social structure (Lukas and Clutton-Brock 2013).

In addition to the capabilities we demonstrated here, `ctmmweb` inherits many of the “under the hood” features of ctmm, including computational algorithms that are highly efficient and scale well to large datasets. The app also has considerably more analytical functionality, including: 1) the ability to accommodate information on telemetry error in all analysis steps, including model fitting/selection and subsequent conditional analyses; 2) scale-free estimation of speed and distance traveled using conditional simulation (Noonan et al. 2019); 3) advanced options for detecting and accounting for multiple sampling schedules in a tracking data; 4) visual subsetting of individual datasets; 5) the ability to detect multiple processor cores and appropriately distribute analyses over them to reduce run time; 6) the ability of perform optimally weighted AKDE home range estimation on irregularly sampled data (Fleming et al. 2018); and 7) occurrence distribution estimation, via time-series kriging (Fleming et al. 2016).

`ctmmweb` is openly available in cloud-hosted form, which requires no installation or configuration, and is also available as a locally installable R package via GitHub, including a wizard-style installer for Windows operating systems. Links to all of these versions, as well as to other `ctmmweb`-related resources can be found in Supplementary Material S2. For first-time users and quick, exploratory analyses, we recommended the cloud-hosted version, while for more thorough analysis or regular use, we recommend installing `ctmmweb` locally. To help improve the app going forward, bug reports, suggestions for improvement, or other feedback can be posted on GitHub.

We will continue to expand and refine `ctmmweb`’s capabilities as the `ctmm` package, and the analytical platform it is based on, continue to develop. Currently, `ctmmweb` is the only point-and-click platform that supports AKDE home range estimation, and it also facilitates many other autocorrelation-explicit analyses. We intend `ctmmweb` to make these sophisticated movement tools accessible to a broader range of ecologists, wildlife professionals, and conservation biologists, without requiring advanced knowledge of R programming.

## ACKNOWLEDGEMENTS

JMC, CHF, MJN, and XD were supported by NSF ABI 1458748, and initial development of `ctmmweb` was supported by this grant as well. JMC, CHF, and XD were also supported by NSF IIBR 1915347. This work was partially funded by the Center of Advanced Systems Understanding (CASUS) which is financed by Germany’s Federal Ministry of Education and Research (BMBF) and by the Saxon Ministry for Science, Culture and Tourism (SMWK) with tax funds on the basis of the budget approved by the Saxon State Parliament.

## SUPPORTING INFORMATION

Additional supporting information may be found in the online version of this article at the publisher’s website.

