## Supplementary Material S1 for "ctmmweb: A graphical user interface for autocorrelation-informed home range estimation"

**SUPPLEMENTARY MATERIAL S1: Session archive and work report**

The full session archive from the jaguar analysis session that produced the results in this article is provided (Supplementary File S1: `ctmmweb_jaguar_session_archive.zip`). This archive includes the work report (`report.html`) that contains additional tables and figures covered in the main text. In addition, the included `plot.zip` archive contains all the figures produced by `ctmmweb` during the analysis session, saved at the resolution indicated at the bottom of the app's sidebar, as well as the data used in the analysis (`combined._data_table.csv`) and cache (`cache.zip`) of the underlying R workspace containing all fitted models and related elements. This cache allows the user, upon loading the session archive into to `ctmmweb`, to pick up where they left off, without first having to re-run all previous analyses.
