## Supplementary Material S2 for "ctmmweb: A graphical user interface for autocorrelation-informed home range estimation"

**SUPPLEMENTARY MATERIAL S2: Online resources for `ctmmweb`**

1) `ctmmweb`'s online documentation, featuring detailed installation instructions, video demonstrations of webapp usage (with subtitles that can be turned on via closed captioning in YouTube), and vignettes on package usage (https://ctmm-initiative.github.io/ctmmwebdoc/).

2) `ctmmweb` GitHub site, featuring installation instructions, source code, and issue reporting (https://github.com/ctmm-initiative/ctmmweb).

3) The clould hosted version of `ctmmweb` (https://ctmm.shinyapps.io/ctmmweb/).

4) The windows all-in-one wizard style installer (https://github.com/ctmm-initiative/ctmmweb/releases/download/v0.2.10/ctmmwebsetup.exe).
