## Supplementary File S1 for "ctmmweb: A graphical user interface for autocorrelation-informed home range estimation": report.html

Work Report of ctmm web-app


### Work Report of ctmm web-app

`[2020-05-07 10:55:58]` App started

```
Package Build Info: 
- version: 0.2.10
- build_date: 2020-05-06
- commit_message: [3ffa3df] 2020-05-06: update text to 0.2.10
```

#### Import

`[2020-05-07 10:55:58]` Recording is On

`[2020-05-07 10:55:58]` Diagnostic info printing to R Console

`[2020-05-07 10:55:58]` Parallel mode enabled

`[2020-05-07 10:56:08]` Using ctmm data

```
jaguar
```

`[2020-05-07 10:56:08]` Data updated

#### Visualization

`[2020-05-07 10:56:09]` Current selected individuals

| identity | start | end | interval (hours) | duration (months) | points | calibrated |
| --- | --- | --- | --- | --- | --- | --- |
| Brutus | 2013-10-19 06:03 | 2014-01-03 05:01 | 1 | 2.57 | 1323 | no |
| Esperanca2 | 2015-04-26 07:01 | 2015-08-30 23:01 | 1 | 4.29 | 2343 | no |
| Teorema | 2013-04-21 17:01 | 2014-01-21 17:01 | 1 | 9.31 | 4860 | no |
| Troncha | 2013-10-22 06:02 | 2014-01-17 17:01 | 1 | 2.96 | 1390 | no |

`[2020-05-07 10:56:09]` saving plot as

```
plot_2_overview_2020-05-07_10-56-09.png
```

`[2020-05-07 10:56:11]` saving plot as

```
plot_5_histogram_2020-05-07_10-56-11.png
```

`[2020-05-07 10:56:12]` Current selected individuals

| identity | start | end | interval (hours) | duration (months) | points | calibrated |
| --- | --- | --- | --- | --- | --- | --- |
| Brutus | 2013-10-19 06:03 | 2014-01-03 05:01 | 1 | 2.57 | 1323 | no |
| Esperanca2 | 2015-04-26 07:01 | 2015-08-30 23:01 | 1 | 4.29 | 2343 | no |
| Teorema | 2013-04-21 17:01 | 2014-01-21 17:01 | 1 | 9.31 | 4860 | no |
| Troncha | 2013-10-22 06:02 | 2014-01-17 17:01 | 1 | 2.96 | 1390 | no |

`[2020-05-07 10:56:12]` saving plot as

```
plot_2_overview_2020-05-07_10-56-12.png
```

`[2020-05-07 10:56:14]` saving plot as

```
plot_5_histogram_2020-05-07_10-56-14.png
```

#### Model Selection

`[2020-05-07 10:56:20]` saving plot as

```
vario_2020-05-07_10-56-20.png
```

`[2020-05-07 10:56:24]` Trying different models…

`[2020-05-07 11:05:59]` Tried Models

| no | identity | model\_type | ΔAICc | DOF mean | DOF area | DOF speed | area (km²) | area CI (km²) | τ[position] (days) | τ[position] CI (days) | τ[velocity] (minutes) | τ[velocity] CI (minutes) | speed (km/day) | speed CI (km/day) | τ |
| --- | --- | --- | --- | --- | --- | --- | --- | --- | --- | --- | --- | --- | --- | --- | --- |
| 1 | Brutus | OUF anisotropic | 0.00 | 9.94 | 16.24 | 608.90 | 355.70 | (204.27 - 548.45) | 4.23 | (2.15 - 8.34) | 23.60 | (20.69 - 26.91) | 23.33 | (22.46 - 24.19) | NA |
| 2 | Brutus | OUF isotropic | 29.68 | 9.12 | 14.84 | 644.04 | 399.12 | (222.59 - 626.33) | 4.66 | (2.26 - 9.6) | 23.93 | (21.04 - 27.21) | 23.33 | (22.46 - 24.19) | NA |
| 3 | Brutus | OU anisotropic | 531.88 | 6.86 | 10.56 | 0.00 | 359.70 | (176.52 - 606.99) | 6.48 | (2.55 - 16.45) | NA | NA | NA | NA | NA |
| 4 | Brutus | OUf anisotropic | 1088.96 | 115.49 | 234.28 | 1167.08 | 214.89 | (188.25 - 243.26) | NA | NA | NA | NA | 29.38 | (28.51 - 30.24) | 14257.31 |
| 5 | Esperanca2 | OUF anisotropic | 0.00 | 31.55 | 57.88 | 109.99 | 41.40 | (31.42 - 52.72) | 2.07 | (1.53 - 2.8) | 4.30 | (3.42 - 5.41) | 26.78 | (24.19 - 29.38) | NA |
| 6 | Esperanca2 | OUF isotropic | 91.42 | 29.83 | 54.62 | 110.23 | 44.74 | (33.67 - 57.37) | 2.19 | (1.6 - 3) | 4.33 | (3.44 - 5.45) | 26.78 | (24.19 - 29.38) | NA |
| 7 | Esperanca2 | OU anisotropic | 291.74 | 28.44 | 51.82 | 0.00 | 41.52 | (30.99 - 53.56) | 2.31 | (1.67 - 3.19) | NA | NA | NA | NA | NA |
| 8 | Esperanca2 | OUf anisotropic | 2889.72 | 335.98 | 707.32 | 2224.20 | 27.57 | (25.58 - 29.64) | NA | NA | NA | NA | 18.14 | (18.14 - 19.01) | 8159.01 |
| 9 | Teorema | OUF anisotropic | 0.00 | 54.24 | 103.43 | 499.61 | 70.79 | (57.81 - 85.07) | 2.57 | (2.07 - 3.2) | 11.63 | (10.34 - 13.08) | 19.01 | (18.14 - 19.87) | NA |
| 10 | Teorema | OUF isotropic | 69.73 | 53.01 | 101.17 | 535.55 | 73.56 | (59.92 - 88.56) | 2.64 | (2.11 - 3.29) | 11.99 | (10.69 - 13.44) | 19.01 | (18.14 - 19.87) | NA |
| 11 | Teorema | OU anisotropic | 1012.14 | 43.95 | 83.33 | 0.00 | 70.59 | (56.25 - 86.54) | 3.20 | (2.51 - 4.09) | NA | NA | NA | NA | NA |
| 12 | Teorema | OUf anisotropic | 4729.02 | 563.66 | 1194.06 | 4712.62 | 47.81 | (45.13 - 50.55) | NA | NA | NA | NA | 18.14 | (18.14 - 19.01) | 10531.00 |
| 13 | Troncha | OUF anisotropic | 0.00 | 14.31 | 24.57 | 307.88 | 168.03 | (108.28 - 240.7) | 3.26 | (1.95 - 5.45) | 16.11 | (13.63 - 19.04) | 22.46 | (20.74 - 23.33) | NA |
| 14 | Troncha | OUF isotropic | 8.77 | 14.68 | 25.36 | 307.99 | 164.34 | (106.72 - 234.2) | 3.17 | (1.92 - 5.25) | 16.13 | (13.65 - 19.07) | 22.46 | (20.74 - 23.33) | NA |
| 15 | Troncha | OU anisotropic | 397.28 | 10.93 | 18.30 | 0.00 | 169.25 | (100.79 - 255.15) | 4.39 | (2.37 - 8.16) | NA | NA | NA | NA | NA |
| 16 | Troncha | OUf anisotropic | 1314.33 | 150.29 | 317.27 | 1328.70 | 103.83 | (92.72 - 115.56) | NA | NA | NA | NA | 24.19 | (24.19 - 25.06) | 11750.45 |

`[2020-05-07 11:06:02]` Selected Models

| model\_no | identity | model\_type |
| --- | --- | --- |
| 1 | Brutus | OUF anisotropic |
| 5 | Esperanca2 | OUF anisotropic |
| 9 | Teorema | OUF anisotropic |
| 13 | Troncha | OUF anisotropic |

`[2020-05-07 11:06:05]` saving plot as

```
vario_2020-05-07_11-06-05.png
```

#### Home Range

`[2020-05-07 11:15:08]` Calculating Home Range in Same Grid …

`[2020-05-07 11:16:00]` Home Range Summary

| no | identity | model\_type | DOF area | DOF bandwidth | quantile | area (km²) | area CI (km²) |
| --- | --- | --- | --- | --- | --- | --- | --- |
| 1 | Brutus | OUF anisotropic | 16.237 | 25.979 | 95 | 296.66 | (170.36 - 457.41) |
| 5 | Esperanca2 | OUF anisotropic | 57.885 | 143.032 | 95 | 37.60 | (28.54 - 47.89) |
| 9 | Teorema | OUF anisotropic | 103.426 | 314.787 | 95 | 61.88 | (50.53 - 74.37) |
| 13 | Troncha | OUF anisotropic | 24.565 | 40.811 | 95 | 144.96 | (93.41 - 207.65) |

`[2020-05-07 11:16:01]` saving plot as

```
home_range_2020-05-07_11-16-01.png
```

`[2020-05-07 11:16:01]` saving plot as

```
home_range_2020-05-07_11-16-01.pdf
```

#### Overlap

`[2020-05-07 11:24:38]` Overlap Summary

| v1 | v2 | CI low | est | CI high |
| --- | --- | --- | --- | --- |
| Brutus | Esperanca2 | 0.26 | 0.43 | 0.62 |
| Brutus | Teorema | 0.51 | 0.71 | 0.88 |
| Brutus | Troncha | 0.24 | 0.43 | 0.66 |
| Esperanca2 | Teorema | 0.17 | 0.23 | 0.31 |
| Esperanca2 | Troncha | 0.38 | 0.57 | 0.76 |
| Teorema | Troncha | 0.09 | 0.18 | 0.32 |

`[2020-05-07 11:24:38]` saving plot as

```
overlap_plot_value_range_2020-05-07_11-24-38.png
```

`[2020-05-07 11:24:46]` saving plot as

```
Overlap of Home Range_2020-05-07_11-24-46.png
```

`[2020-05-07 11:24:47]` saving plot as

```
Overlap of Home Range_2020-05-07_11-24-47.pdf
```

#### Map

`[2020-05-07 11:24:56]` Saving map: Point

Point

`[2020-05-07 11:25:11]` Saving Data

`[2020-05-07 11:25:11]` All Telemetry Data

| identity | start | end | interval (hours) | duration (months) | points | calibrated |
| --- | --- | --- | --- | --- | --- | --- |
| Brutus | 2013-10-19 06:03 | 2014-01-03 05:01 | 1 | 2.57 | 1323 | no |
| Esperanca2 | 2015-04-26 07:01 | 2015-08-30 23:01 | 1 | 4.29 | 2343 | no |
| Teorema | 2013-04-21 17:01 | 2014-01-21 17:01 | 1 | 9.31 | 4860 | no |
| Troncha | 2013-10-22 06:02 | 2014-01-17 17:01 | 1 | 2.96 | 1390 | no |

`[2020-05-07 11:25:11]` Work Report Generated
